# Denoising image-based spatial transcriptomics data with DenoIST

**DOI:** 10.1101/2025.11.13.688387

**Authors:** Aaron Wing Cheung Kwok, Annika Vannan, Nicholas E. Banovich, Jonathan A. Kropski, Heejung Shim, Davis J. McCarthy

**Author notes:** Contributing authors.

## Abstract

Image-based spatial transcriptomics (IST) technologies provide unprecedented resolution of gene expression in tissue sections, but suffer from contamination of cells’ gene expression profiles due to imperfect cell segmentation. We present DenoIST (Denoising Image-based Spatial Transcriptomics data), a new computational tool that accurately identifies and removes contaminating transcripts from IST datasets. DenoIST models the observed transcript counts using a Poisson mixture model that explicitly accounts for local neighbourhood contamination. Applied to multiple real IST datasets of varying cell densities, DenoIST restores gene expression specificity and clarifies local biological structures by identifying and filtering transcripts spilt over from neighbouring cells. The denoised data enable more consistent and interpretable cell type annotation by minimising conflicting gene expression profiles, reducing the prevalence of hybrid or ambiguous cell types, and enhancing the contrast between distinct functional compartments. Overall, we demonstrate that DenoIST can be integrated to existing IST analysis workflows to improve biological interpretability and robustness of IST data.

## 1 Introduction

Spatial transcriptomics (ST) has emerged to be one of the most popular assays in recent years for studying the localisation of biological activities and pathogenic processes. As always, with new assays come new bioinformatics challenges, and the field is still looking for best practices for various analysis tasks. Initial iterations of ST were mostly based on Next Generation Sequencing (NGS), such as 10x Visium [1], which motivated the application of computational methods designed for single cell RNA-sequencing (scRNA-seq) data as they share similarities in both the biology of interest and NGS-based assaying techniques. However, recent developments in image-based spatial transcriptomic (IST) assays, such as Vizgen MERSCOPE [2], NanoString CosMx [3], and 10x Genomics Xenium [4], enabled the possibility of identifying individual transcripts at sub-micron resolution using optical microscopy approaches. These new technologies presents new bioinformatics challenges that are not trivial to resolve using existing tools for scRNA-seq data analysis, as the data generative processes and data characteristics differ greatly from those familiar from NGS-based assays.

One of the major unresolved challenges is the ‘transcript contamination’ problem of IST data. ‘Transcript contamination’ refers to when gene transcripts from one cell are erroneously assigned to nearby cells due to segmentation errors, causing some cells to present gene expression profiles that are biologically implausible. This problem is a manifestation of the broader challenge that is cell segmentation, i.e., the inference of cell boundaries in the assayed tissue. Unlike in single-cell assays where sequencing libraries can be constructed in which transcripts from each single cell can be identified by barcode labels, IST relies on computational approaches that may use staining of cell nuclei and/or membranes, transcript density, or other available information, to define cell boundaries and assign transcripts into the inferred partitions. Unfortunately, even in theory, perfect cell segmentation is almost impossible if only the 2D planar projection is considered while in reality the cells exist in a 3D space; causing overlapping events on the vertical axis to be missed [5]. Fully utilising the z-axis in current datasets is technically challenging because microscopic resolution is lower along the vertical dimension [5]. Despite the field’s best efforts in advancing cell segmentation methods [6–9], the limitations of our physical reality are hard to overcome.

As such, ‘transcript contamination’ is a pervasive problem in IST analysis. Apart from the obvious fact of adding unwanted noise to observed cellular gene expression profiles, transcript contamination has two main implications for downstream analysis: 1) hybrid cell types will be ‘found’ when it is likely to be contamination instead of true biology; and 2) varying degrees of contamination will confound quantitative comparisons between samples. These issues are direct consequences of limitations in cell segmentation with extensive impact on routine tasks such as differential expression analysis [10].

Given imperfect cell segmentation, there are two major ways to mitigate the transcript contamination issue. First, one might limit analyses to transcripts within a cellular feature for which segmentation is much more reliable than it is for whole cells. The most obvious and common approach of this kind is to analyse transcripts assigned to well-segmented cell nuclei only. These transcripts are typically of high confidence and present with fewer contamination effects. Although segmentation of nuclei typically cannot avoid contamination caused by overlapping of cell/nuclei in the Z dimension, staining for cell nuclei is generally much more trustworthy than inferred boundaries of whole cells. However, Vannan et al. [11] observed that even with this conservative approach, in which we may only retain 10% of a cell’s transcripts [12], contamination remains a significant problem. As such, this approach is often unacceptably conservative and may discard the vast majority of observed transcripts (and with them important biological signal) without solving the contamination issue sufficiently to avoid confounding downstream analyses.

The second way to mitigate contamination effects is to apply count correction methods to denoise the data prior to analysis. At the time of writing, there are two methods that are designed for this purpose: resolVI [13] and SPLIT [14]. ResolVI is a method extended from the scVI model [15], which uses a variational autoencoder (VAE) to learn the latent representation of spatial transcriptomics data. To account for marker contamination, the decoder model in resolVI parameterises the contamination effect as latent variables and can return ‘corrected counts’ by sampling from the corrected latent embeddings. Alternatively, SPLIT is a method that uses an scRNA-seq reference to determine whether cells in IST are “pure” or a mixture of cell types. Briefly, it uses RCTD [16], a method for deconvoluting cell-type proportions in sequencing-based spatial data, to learn from the reference data whether each cell is a doublet of two different cell types and the fraction of transcripts from each cell type. To correct the counts, the raw counts are weighted by the expected fraction of transcripts from the primary cell type.

Both denoising methods, resolVI and SPLIT, have their limitations. First, resolVI does not ‘correct’ counts by modifying the raw counts, instead it samples from the latent space a new set of ‘counts’ that are corrected for the effects specified in the VAE. These imputed counts do not exist on the spatial level, which might affect one’s interpretation of spatial neighbourhoods. Second, SPLIT requires an annotated scRNA-seq reference that is not always available for specific tissues and/or disease contexts. Lastly, both tools output non-integer corrected counts, which complicate interpretation.

To address this research gap, we present DenoIST (Denoising Image-based Spatial Transcriptomics), a Poisson mixture model tailored for denoising IST data by reducing the effects of transcript contamination in downstream analysis tasks. Importantly, DenoIST requires no prior cell-type information or reference data and returns integer counts. We demonstrate its application on various IST datasets and show that DenoIST is effective in contamination removal and improves downstream analysis.

## 2 Results

### 2.1 Segmentation artefacts in IST data introduces contamination and spurious findings

Cell segmentation for IST data is an unresolved challenge and artefacts arising from mis-segmentation propagates to downstream analysis in various ways [10]. Here we further illustrate the issues discussed by Vannan et al. [11] in their lung fibrosis study. Cells in the lung were highly challenging to segment accurately due to the irregular structures of the alveoli and the blood vessels interweaving between air spaces. As a result, segmentation errors were prevalent as biologically implausible gene profiles were frequently observed in the count data. A visual example is shown in Fig. 1a using one of the healthy lung samples. Even with a better cell segmentation method, markers for the myeloid lineage are often found together with that of the endothelial, epithelial, and alveolar lineages. Co-expression of these supposedly mutually exclusive lineage markers suggests a high degree of contamination due to segmentation limitations.

**Fig. 1.**
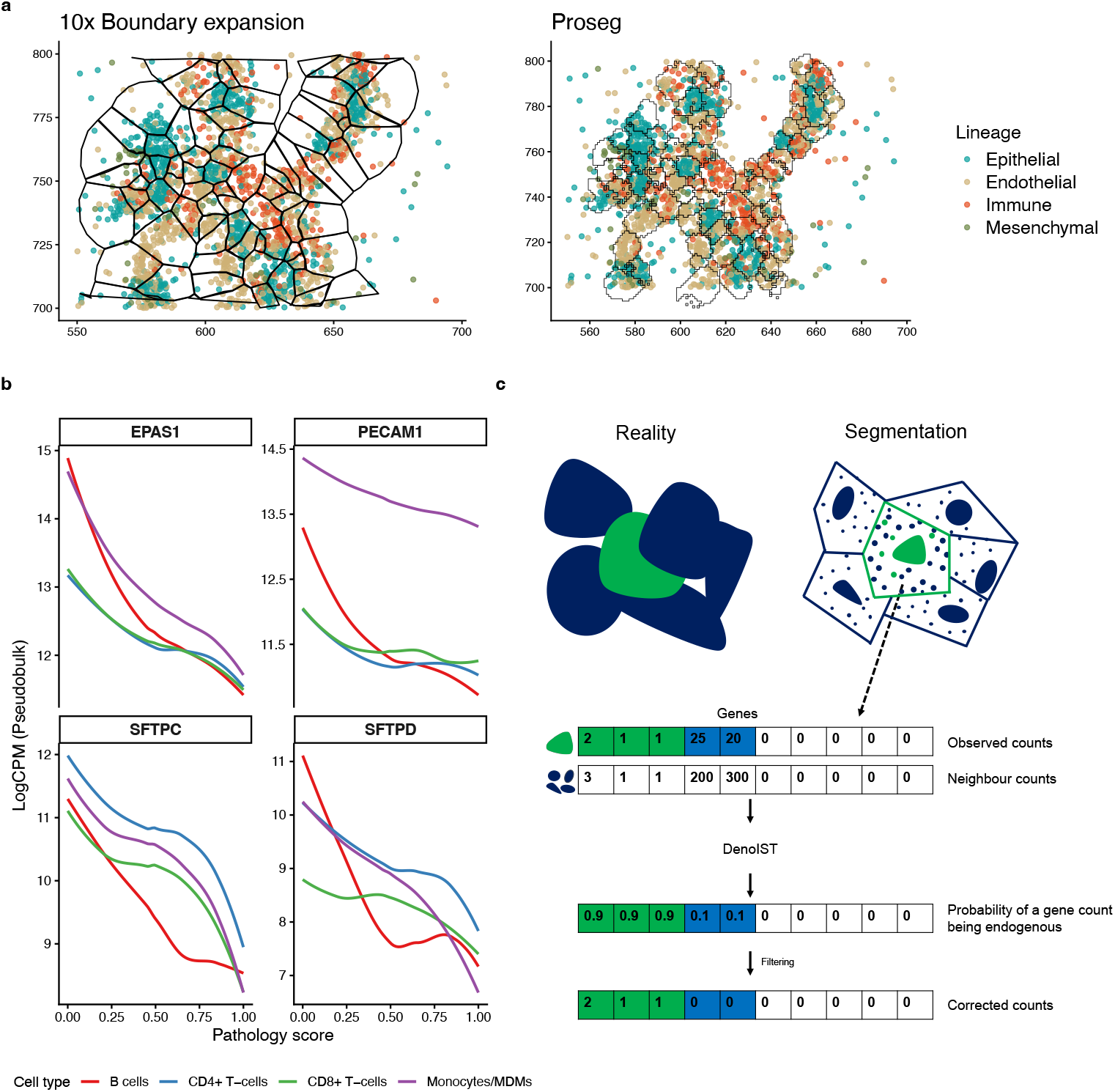
a) Shown here is a region of the healthy lung sample assayed using Xenium, with the boundary expansion segmentation from 10x Xenium Ranger (Top) and Proseg (Bottom) segmentation. Each dot is a transcript molecule, molecules of lineage marker genes are coloured by their respective lineages. Black lines are segmentation boundaries. b) Log CPM (counts per million) normalised pseudobulked gene expression of 4 selected contaminating genes using 4 annotated immune cell types in the lung fibrosis Xenium data. Each data point is a sample and a lowess curve is fitted over all the samples. c) A high level schematic of the application of DenoIST.

The contamination complicates downstream analysis tasks, for example the varying degrees of contamination between samples can confound real biological differences. This confounding can be observed when trying to compare gene expression between healthy and diseased samples. Using the pseudobulked counts from only the annotated immune cells, genes that are not expected to be expressed in immune cells are found to be highly enriched in healthier samples over diseased ones (Fig. 1b). Most of these genes are either endothelial markers (*EPAS1, PECAM1*) or alveolar markers (*SFTPC, SFTPD*). This observation is due to healthy samples typically having more vasculature than diseased samples, leading to more contamination from endothelial cells for healthier samples, and systematically more endothelial transcripts in these immune cells.

### 2.2 Overview of DenoIST model

DenoIST is a computational method for detecting contaminated genes in each segmented cell in IST count data. The input to DenoIST is a gene × cell count matrix generated from any IST assay (e.g., Xenium, MERSCOPE, CosMx, etc.) with any cell segmentation algorithm, along with the spatial coordinates of each cell. The output is an adjusted count matrix with the contaminated gene counts removed (Fig. 1).

The DenoIST model works as follows. Briefly, each cell is assumed to have a proportion of genes that are truly expressed due to biology. The rest of the genes are not expressed, but can have non-zero counts due to contamination, that is, incorrect assignment of non-endogenous transcripts to the cell. With this mental model, we can formulate tackling contamination into a mixture model problem and infer which gene counts are truly endogenous for each cell. We can then remove the gene counts from genes that are inferred to be contaminated.

DenoIST considers a cell’s neighbourhood counts to determine if a gene count is endogenous. If a segmented cell is taken out of its spatial context, then the higher the gene count, the more likely it is endogenously expressed. However, in spatial data these cells do not exist in a vacuum. The absolute gene count of a cell should be weighed against the cell’s local neighbourhood to make sense. In an illustrated example (Fig. 1c), 25 transcripts for a gene in a cell are a lot if we consider this cell alone, but if its local neighbourhood is flooded with the same transcripts, then the 25 transcripts are no longer as important as before. Conversely, 2 transcript counts for a gene can be very important if they are the only 2 transcripts in its local neighbourhood.

This principle underlies how DenoIST distinguishes endogenous expression from contamination. By comparing each cell’s gene counts against those of its local neighbourhood, DenoIST identifies transcripts that are disproportionately abundant in nearby cells relative to the target cell’s own profile. Such transcripts are likely contaminants originating from segmentation inaccuracies, diffusion, ambient contamination, or other unknown processes rather than true endogenous expression. Conversely, transcripts whose counts remain high relative to the local background are more likely to be endogenously expressed. Through this neighbourhood-aware framework, DenoIST effectively suppresses contamination while preserving genuine biological signal.

### 2.3 DenoIST removes contamination from Xenium human breast cancer data

We first performed a case study on the Xenium human breast cancer dataset [4] by applying DenoIST on the dataset with cells segmented using the boundary expansion algorithm from Xenium Ranger. To evaluate DenoIST’s performance on noise removal, we first checked whether marker genes are correctly localised to known structures. One example is *ACTA2*, which is a marker gene for smooth muscle cells and in particular marks the basal zone in DCIS [17]. With the default segmentation, *ACTA2* is persistently detected in cells nearby the breast ducts, either due to mis-assignment or diffusion. DenoIST correctly identifies cells that are contaminated and the denoised data shows high specificity of *ACTA2* in the duct walls (Fig. 2a), which corroborates with immunohistochemistry staining results [17].

**Fig. 2.**
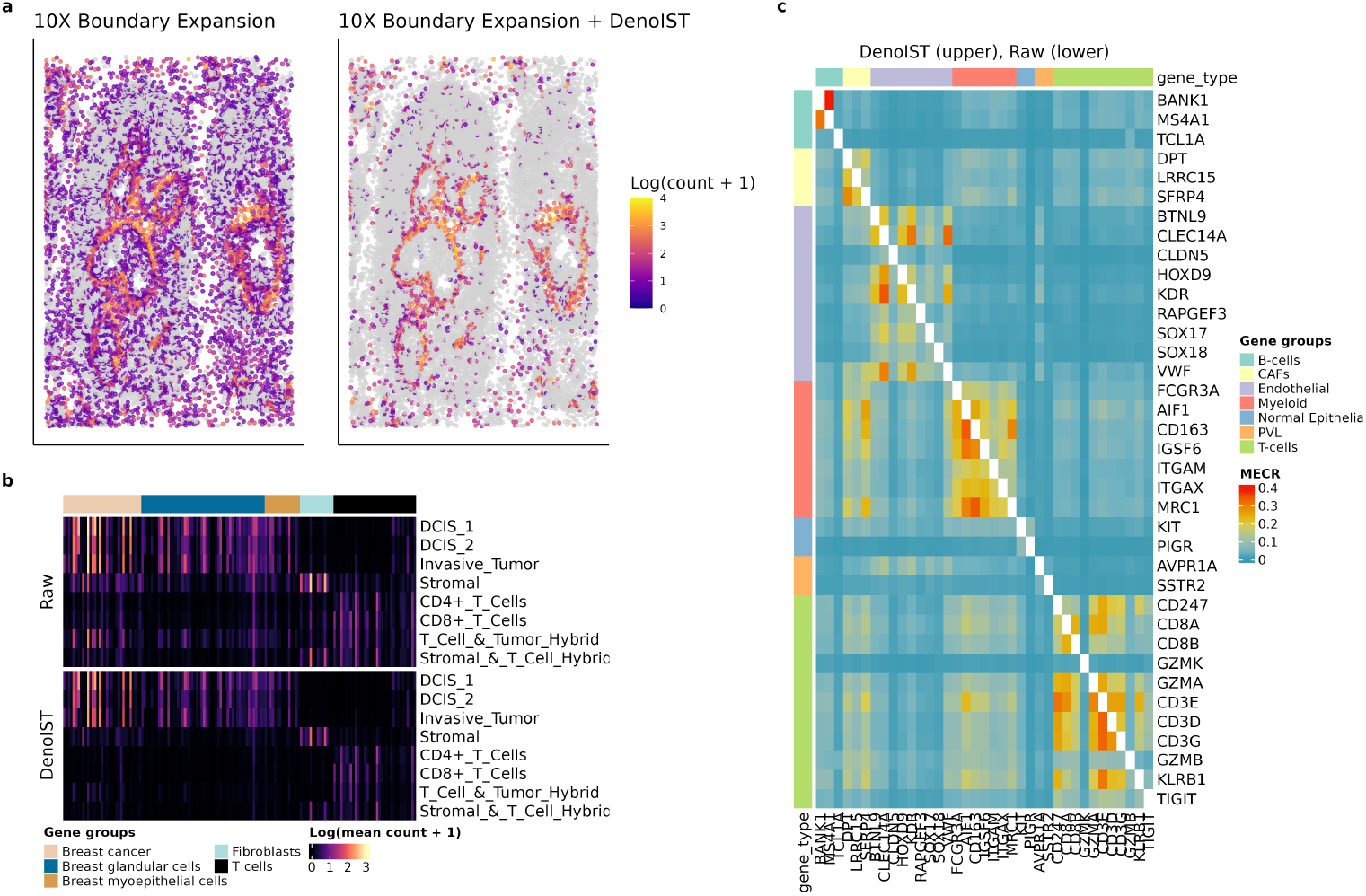
Expression of *ACTA2* from a zoomed-in section in Xenium human breast cancer dataset. Each dot is a segmented cell using 10x boundary expansion method, colour shows the log count of *ACTA2*. Cells with 0 count are greyed out for visual clarity. b) Heatmap visualisation of gene expression of annotated cell types before (top) and after DenoIST (bottom). Columns are selected genes, annotated by the cell type they mark. Row are cell types. The log(mean count + 1) for each cell type is shown here. c) MECR before (top) and after (bottom) applying DenoIST to Xenium human breast cancer dataset. Rows and columns denote genes and each entry is the MECR of the corresponding pair. Note that genes that mark the same cell type are not expected to be mutually exclusive, but are shown here for positive control.

DenoIST-adjusted data also shows there is a lack of evidence for novel hybrid cell types. We compared the gene group annotations with the cell type annotations (both provided by 10x Genomics), and observed higher correspondence between the annotations after applying DenoIST (Fig. 2b). For example, annotated T cells were found to also express many breast cancer and breast myoepithelial genes in the raw counts, but DenoIST identified these improbable cases as contamination. Moreover, there are cell populations that were annotated as hybrids because they highly express genes from several gene groups. However, after applying DenoIST, these hybrid cell types no longer have conflicting gene profiles and resolve to purer cell types. For example, the gene expression profile of T Cell and Tumour hybrid resembles that of T cells after denoising, instead of ubiquitously expressing cancer and T cell gene markers.

To systematically evaluate the improvement of denoised counts, Mutually Exclusive Coexpression Rate (MECR) was used to quantify the reduction in false positives (see Methods in Section 5.7.1). Briefly, MECR measures the co-expression rate of gene pairs that are known to be mutually exclusive. As these gene pairs are mutually exclusive in single cell reference datasets, their co-expression in spatial datasets must be a technical artefact. Hence, a good denoising method should yield data with low MECR.

Analysing MECR of 337 mutually exclusive gene pairs derived from a single cell breast cancer atlas from Wu et al. [18] provides compelling evidence of false-positive transcript assignments to cells (Fig. 2c). Boundary expansion is a relatively lax segmentation method and often results in transcript mis-assignment, which is reflected by the high MECR metric. After running DenoIST, MECR is substantially reduced while preserving co-expression patterns between genes that mark the same cell types.

### 2.4 DenoIST outperforms similar tools

DenoIST was benchmarked against two other count correction tools, resolVI [13] and SPLIT [14], using five IST datasets with varying tissue densities assayed with different technologies: Xenium, MERSCOPE, and CosMx mouse brains (most dense), Xenium human breast cancer, Xenium human healthy lung (least dense) (Table 1); and three different segmentation methods: default (most lenient), 10x nuclei (most conservative), and Proseg (most sophisticated).

**Table 1.** Summary of datasets used.

| Dataset | Assay | Citation | Accession | # cells | # genes | Ref. dataset |
| --- | --- | --- | --- | --- | --- | --- |
| Human breast cancer | Xenium | Janesick et al. [4] | 10x website | 166,363 | 313 | Wu et al. [18] |
| Mouse brain | Xenium | 10x Genomics | 10x website | 162,033 | 248 | Zeisel et al. [21] |
| Human lung fibrosis | Xenium | Vannan et al. [11] | GSE25036 | 1,630,319 | 343 | Natri et al. [22] |
| Mouse brain | MERSCOPE | Vizgen | Vizgen website | 78,329 | 483 | Zeisel et al. [21] |
| Mouse brain | CosMx | Nanostring | Nanostring website | 48,180 | 950 | Zeisel et al. [21] |

DenoIST is highly effective in reducing MECR, and it is consistent throughout multiple datasets and varying qualities of cell segmentation (Fig. 3a). Comparing to resolVI and SPLIT, DenoIST is largely agnostic to the initial segmentation quality and provides consistent improvement. It outperforms resolVI and SPLIT in most scenarios, but the tools’ differences diminish when a better segmentation algorithm is used, i.e., the improvement over unadjusted data is smaller for Proseg than default segmentation. This observation is reasonable as well segmented data should have less noise to begin with.

**Fig. 3.**
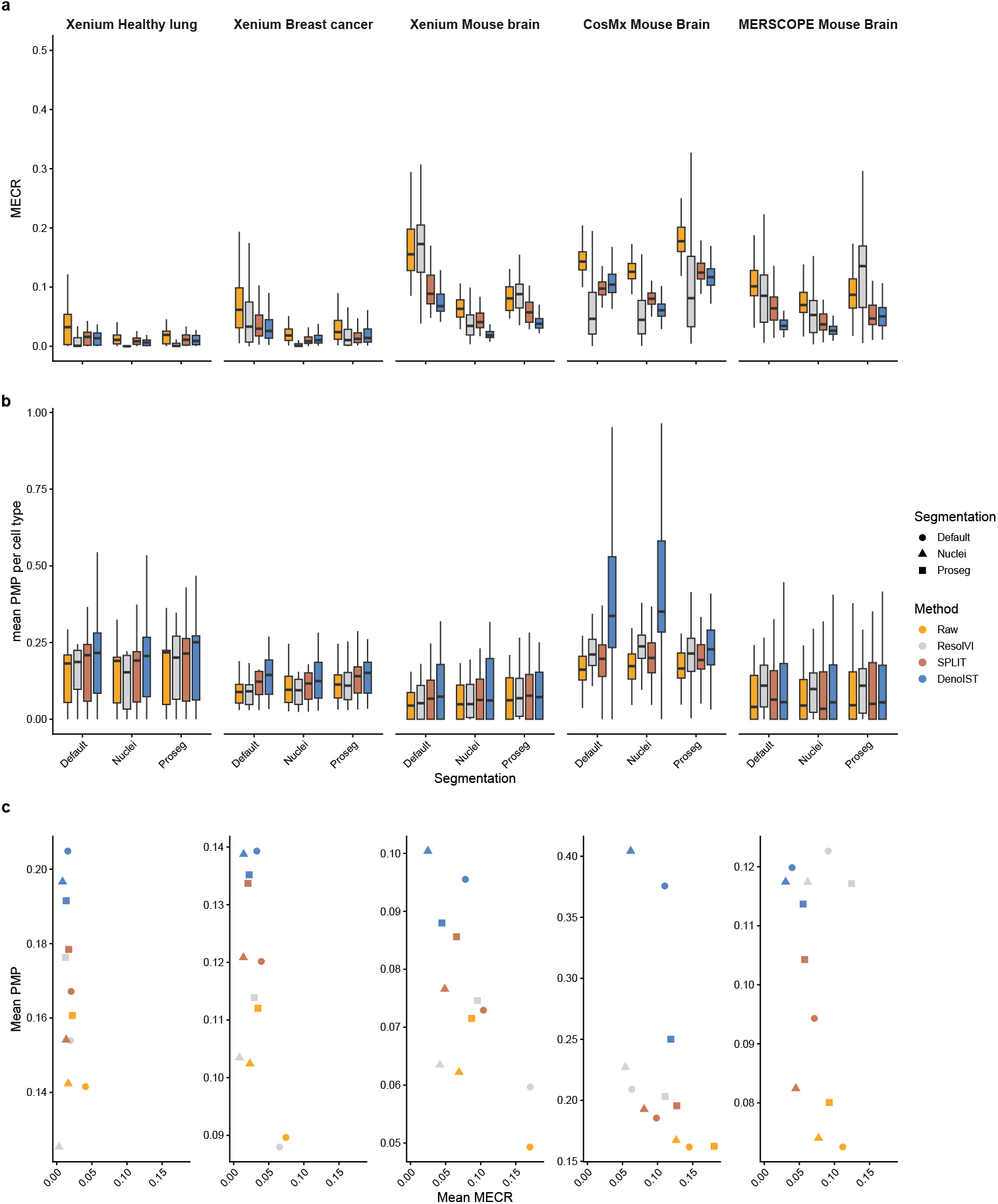
Benchmarking performances of count correction tools across datasets and segmentation methods. a) Comparison of performance on MECR over multiple datasets. Each boxplot denotes the MECR of all the tested mutually exclusive gene pairs. b) Comparison of performance on mean PMP over multiple datasets. Each data point is the mean PMP of a cell type. c) Mean PMP plot against mean MECR shows the trade-off between true and false positives over multiple datasets, segmentation algorithms, and count correction methods.

The MECR metric can be somewhat cheated as hyper-aggressive removal of transcripts without differentiating between true signal and contamination can still lead to a substantial reduction in false positives. Therefore, the true positives should also be evaluated. For this task, I used the Positive Marker Purity (PMP) metric (see methods in Section 5.7.2) proposed by Heidari et al. [6]. Briefly, PMP measures the proportion of positive markers in each cell, with the ‘positive’ markers being defined from an scRNA-seq reference. As PMP serves as a true positive metric, a good denoising method should also yield data with high PMP.

DenoIST achieves satisfactory mean PMP over all tested datasets, again showing highly consistent performance (Figure 3b). Similar to MECR, DenoIST outperforms ResolVI and SPLIT in the true positive metric, showing that the decrease in false positive rate is not due to excessive removal of counts. It is worth noting that resolVI does not show significant improvement over unadjusted data in terms of true positives even though a large reduction in false positives is observed. In some cases, it even has slightly worse PMP than the unadjusted data. This result is somewhat surprising, but consistent with the claim from Bilous et al. [14] that resolVI-corrected data significantly lowers the number of genes expressed and risks over-correction.

Visualising MECR and PMP together emphasises the trade-off between true and false positives (Fig. 3c). It is worth noting that the difficulty of denoising data is related to the biological tissue. For example, in a sparse sample like a healthy lung, all methods achieve relatively low false positives, while in a dense tissue like the mouse brain, contamination and mis-segmentation are harder to avoid, resulting in more false positives and less true positives. However, relative to resolVI and SPLIT, DenoIST provides a good overall balance between MECR and PMP, regardless of dataset and segmentation method.

To demonstrate DenoIST’s capability and consistency on IST assays other than Xenium, we also applied it to MERSCOPE and CosMx mouse brain data. Hartman and Satija [19] previously noted that the different aggressiveness of segmentation algorithms used by 10x Genomics and Vizgen drives between-dataset MECR differences, as a more conservative segmentation should naturally result in a lower false positive rate. This difference is akin to the comparison between 10x nuclei and 10x boundary expansion. Here, we expanded the comparison to CosMx and also evaluated PMP, the true positive metric. Our results show that DenoIST is applicable to other IST assays such as MERSCOPE and CosMx, reflected by similar improvements in both MECR and PMP metrics (Fig. 3d,e). Notably, the base MECR for MERSCOPE data is substantially lower than for the other two assays. This observation is consistent with findings from Hartman and Satija [19]. However, even with a lower base contamination rate, applying DenoIST still led to noticeable improvement in metrics.

### 2.5 DenoIST can be integrated into existing analysis frameworks and provide better biological insights

To demonstrate DenoIST on a practical analysis task, we re-analysed the human lung fibrosis data from Vannan et al. [11]. The original analyses were restricted to nuclei transcripts due to the contamination issue, but some degree of contamination still remained and required post-hoc filtering in downstream analyses [11]. Challenging cell segmentation is unfortunately a limitation of the lung tissue, as its cells have non-typical shapes and the 3D structure of alveoli is hard to capture accurately. We expect a proper denoising tool to mitigate the issue of transcript mis-assignment and perhaps boost signal in downstream tasks.

Studying two example genes before and after running DenoIST neatly distils the broader effects of applying DenoIST (Figure 4a). *PECAM1* and *COL1A1* are gene markers for endothelial and mesenchymal lineages respectively, however they are found in many other lineages due to contamination, even when only analysing nuclei transcripts. After running DenoIST, the non-specific signal is mostly removed, restoring specificity to their respective lineages. Although the data is restricted to nuclei transcripts only, low ambient noise is still prevalent due to the sheer abundance of some specific cell types. Across these six example genes, DenoIST significantly reduces contamination from lineage specific genes while preserving the sensitivity for the correct cell type, without using cell-type information as a prior.

**Fig. 4.**
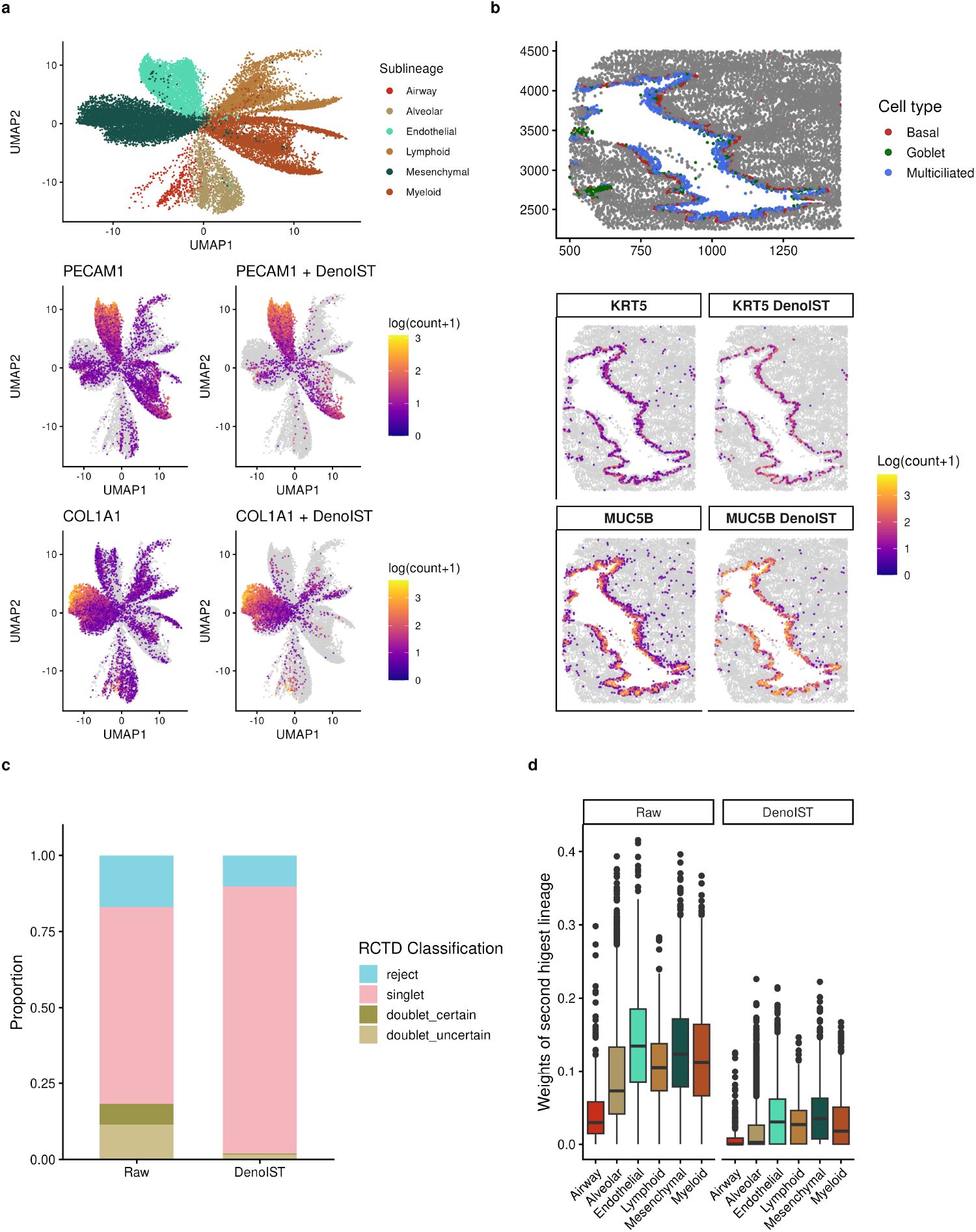
a) UMAP visualisation of lung fibrosis data after applying DenoIST. Sample TILD028MA is shown here. Cells with 0 count are greyed out for visual clarity. b) An airway section from fibrotic sample VUILD110. Each dot is a cell. Cells with 0 count are greyed out for visual clarity. Annotated airway cell types and gene expression (raw counts and DenoIST-adjusted counts) for *KRT5* and *MUC5B* are shown. c) Proportions of RCTD classification using raw counts and DenoIST-adjusted counts in healthy sample VUHD116A. d) RCTD assignment weights of the second highest lineage of each cell in healthy sample VUHD116A, stratified by their manually annotated lineages. Cells with a pure identity should have low weights for the incorrect lineages.

On the spatial level, data processed with DenoIST can more faithfully capture local structure. For example, the lung airway should exhibit a layered structure as multiciliated cells and goblet cells should sit on top of basal cells [20]. However, this structure is not accurately reflected by their respective marker genes (e.g., *KRT5* for basal cells, *MUC5B* for multiciliated and goblet cells) as they often ‘bleed’ through layers due to contamination from nearby cells (Fig. 4b). However, DenoIST correctly identifies contamination and the resulting gene expression resembles a more clearly defined layer structure. This improvement in delineation of fine-scale spatial structure shows the importance of denoising IST data as more nuanced biological spatial relationships can be masked by mis-segmentation and contamination.

We further show that DenoIST reduces conflicting gene expression profile by applying RCTD to the lung data. RCTD was originally designed for deconvoluting spot-based spatial data where each spot is large enough to accommodate multiple cells [16]. We expect in IST where subcellular resolution data is available, most segmented cells should have a pure identity and ‘doublets’ should be infrequent. However, this is not the case in the lung data as RCTD identified only 64.8% of cells as ‘singlets’, with the rest being confident doublets or having ambiguous identities (Fig. 4c). The denoised data showed a higher proportion of singlet classifications (87.9%) compared to the raw data, indicating that DenoIST reduced signal ambiguity across segmented cells. In addition, the second-highest cell type weight for each cell decreased after denoising, suggesting greater separation between the top and secondary assignments (Fig. 4d). Together, these results demonstrate that DenoIST enhances the reliability of cell type assignment in IST data.

## 3 Discussion

In this study, we developed DenoIST and demonstrated its capability in denoising IST data and versatility in different biological and technical settings. Importantly, DenoIST outperforms similar tools in both false positve and true positive metrics, while requiring minimal inputs or supervision. Compared to resolVI, DenoIST does not over-correct and is less computationally expensive; compared to SPLIT, DenoIST does not require a scRNA-seq reference. With its transcript filtering approach, DenoIST returns integer counts unlike resolVI and SPLIT, which instead return weighted non-integer counts that might be hard to interpret as these fractional values do not correspond to a transcript present in the spatial assay. DenoIST is also compatible with all segmentation algorithms and does not over-correct when the segmentation is of high quality, showing its utility on well-segmented data without having the concern of losing real biological signal.

DenoIST makes two key assumptions of which users should be aware. First, it assumes gene counts are all-or-nothing, i.e., genes are either exclusively truly expressed or contaminated. In reality, this assumption over-generalises how transcript counts are quantified, as any given gene can potentially have some transcripts that are ‘real’ (endogenous) and some that are ‘contamination’. Ideally, each transcript should be considered case-by-case instead. However, on datasets with relatively small and targeted gene panels, genes are usually selected to be highly specific to the biology of interest, with most of them being cell-type specific markers. Measuring house keeping genes that are ubiquitously expressed in all cell types is usually a waste of budget. This experimental design implies most genes in the panel should be mutually exclusive of each other to some extent, which explains why DenoIST is effective with this bold assumption that seems demonstrably untrue. However, when the gene panel extends to the whole transcriptome, i.e., when the gene panel size of IST catches up to that of sequencing-based assays, e.g., the 5,000 gene panel in Xenium Prime, this assumption might not hold and needs to be reconsidered. This limitation is therefore especially relevant for future IST datasets with much larger gene panels.

The second assumption is that the truly expressed genes in each cell share the same Poisson mean and the expression level is unimodal. In reality, these assumptions are unlikely to be strictly true as genes generally exhibit a wide dynamic range of expression levels (e.g., as assayed with scRNA-seq), such that they do not share the same Poisson mean. Even within a cell, some genes can be more strongly expressed than others, depending on the genes’ characteristics and functional importance. However, in the context of learning which genes are ‘contamination’, the difference between true expression and contamination is large enough for this assumption to work empirically. Similar to the first assumption, for a large gene panel with more nuances in gene expression, this assumption may negatively effect denoising performance. Future development of the model should aim to account for biological variation between genes in the same cell to establish a more accurate per-cell model. However, as current IST technology tends to produce data with low sensitivity and low dynamic ranges in gene count, DenoIST is applicable in this particular setting.

The major limitation of DenoIST is scalability, as a separate model is fit per cell. In IST datasets with a large number of cells, model fitting can be a bottleneck. While this issue can be partially solved via providing more CPUs and memory to parallelise over, there is room for better optimisation in the model fitting procedure by utilising more low level programming languages. Moreover, if some cell-type information is available, cells of the same cell type should express similar sets of genes. Information sharing within cell types can speed up the model fitting step and perhaps even improve the fit, however this possibility has not been explored here.

Another limitation is that the varying count adjustment accuracy of different samples can potentially introduce more batch effects. We observed that DenoIST is less effective in correcting the healthy lung samples compared to the diseased ones. This difference is likely due to healthy lung samples being too structurally sparse and the local neighbourhood information not providing enough evidence for identifying contamination. As a result of different degrees of adjustment, DenoIST might introduce more batch effects into multi-sample datasets like the lung dataset used here. However, this potential pitfall is not explored due to time and scope constraint.

## 4 Conclusion

To conclude, we have developed DenoIST, an R package for denoising IST data. We have shown that DenoIST works in a range of biological settings and is applicable to most IST assays. We have also demonstrated how DenoIST can be used to improve existing IST analysis workflows and enhance signal purity. Although segmentation-agnostic analysis methods exist, it is often more intuitive to ask questions based on the fundamental unit of biology, which is the cell. Therefore, we expect count correction or denoising to be a key step in future IST data analysis workflows.

## 5 Methods

### 5.1 Datasets

IST datasets and reference scRNA-seq datasets used in this study are listed in Table 1. Xenium human breast cancer dataset was downloaded from https://www.10xgenomics.com/products/xenium-in-situ/preview-dataset-human-breast, only replicate 1 was used. Xenium mouse brain dataset was downloaded from https://www.10xgenomics.com/datasets/fresh-frozen-mouse-brain-replicates-1-standard, only replicate 1 was used. MERSCOPE mouse brain dataset was downloaded from https://info.vizgen.com/mouse-brain-data, only slide 1 replicate 1 was used. CosMx mouse brain dataset was downloaded from https://nanostring.com/products/cosmx-spatial-molecular-imager/ffpe-dataset/cosmx-smi-mouse-brain-ffpe-dataset/, the hemisphere data (slide 2) was used.

### 5.2 DenoIST model

To solve the transcript contamination problem, we propose a model based on the data generative process of IST data (Supplementary Note 1). We work with an *N* × *G* ‘count matrix’ **X** which holds information about the number of observed counts in *N* cells and *G* genes. We let *i* ∈ {1, …, *N* } index cells and *j* ∈ {1, … *J*} index genes, so that *x*_*ij*_ is the observed count of the *j*th gene in the *i*th cell. Then, the observed counts of each cell is modelled as a mixture of endogenously expressed genes and unexpressed (i.e., non-endogenously expressed) genes. Unexpressed genes include zero count genes that are not observed, and non-zero count genes that are only observed due to contamination:

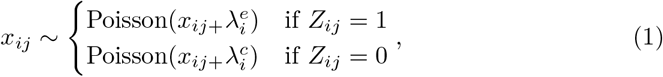

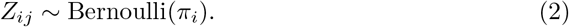

Where:

*π*_*i*_ denotes the proportion of genes that are endogenously expressed in cell *i*,

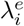 denotes the mean parameter of genes that are endogenously expressed in cell *i*, and 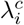 denotes the mean parameter of genes that are not expressed in cell *i*. Note that if there is no contamination or error in signal detection, 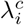 is expected to be 0.

Lastly, *x*_*ij*+_ is defined as the sum of local and global contexts, which we explain below.

#### 5.2.1 Local context

The offset term *x*_*ij*+_ is calculated by:

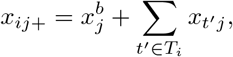

where *T*_*i*_ denotes the set of neighbouring cells to cell *i*, and 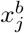 the global background offset to account for ambient contamination. The default option to define *T*_*i*_ is to include cells with centroids within 50 µm from the centroid of cell *i*. A distance based instead of an N-nearest cells based inclusion is to account for neighbourhood densities, as denser tissues tend to exhibit more contamination due to cells being more closely packed together and segmentation difficulty is higher. However, this distance *d* is tunable by the user.

#### 5.2.2 Global context

To account for global ambient contamination over the whole sample, a background count 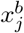 that is shared globally among all cells can be added to *x*_*ij*+_. To calculate 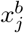, transcripts are binned into 200 hexbins by default. To determine which spatial bins represent background (i.e., areas not primarily occupied by cells), a Gaussian Mixture Model (GMM) is used. The total transcript counts across all spatial bins are assumed to follow a 2-component mixture of Gaussian distributions: one representing background with low transcript counts and the other corresponding to bins with biologically meaningful signal, i.e., higher transcript counts.

Formally, let *C*_*k*_ denote the total transcript count in hexbin *k*. The GMM models the density of *C*_*k*_ as:

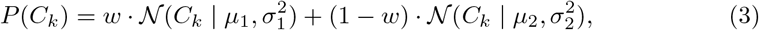

where *w* denotes the proportion of background bins, *µ* and *σ*^2^ are the mean and variance of their respective components. The GMM is fit using the Expectation Maximisation (EM) algorithm, as implemented in the ‘flexmix’ R package. After model fitting, each hexbin is assigned to the most likely component based on its posterior probability. The component with the lower *µ* is designated as the background component. The global background expression level for gene *j*, denoted 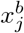, is then computed by averaging the gene’s expression values across all bins classified as background, and then scaled to the local neighbourhood size defined by the distance *d*:

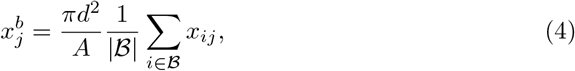

where ℬ denotes the set of bins assigned to the background component, and *A* denotes the area of one hexbin which can be calculated with basic geometry, based on the size of the input sample and number of bins used.

This global background profile is subsequently incorporated into the DenoIST model as an additive offset to account for both local and global context.

### 5.3 Parameter estimation

The EM algorithm is used to estimate parameters for the model. Briefly, the EM algorithm is an iterative method for finding maximum likelihood estimates of parameters by alternating between estimating the expected values of the latent variables (E-step) and maximising the likelihood with respect to the parameters (M-step) [23]. The set of parameters needed to be estimated from data is 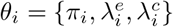. Detailed derivations are described in Supplementary Note 3.

Multiple initialisation is used to ensure robustness. Briefly, 10 initial values for *π*_*i*_ are drawn from Unif(0, 0.5), then the fit with the highest log likelihood is chosen as the best fit. The rationale of capping the initialisation value of *π* to 0.5 is based on the intuition that a cell is unlikely to truly express more than 50% of the genes in the gene panel, given that most gene panels in currently available IST assays are targeted panels instead of whole transcriptome.

### 5.4 Posterior probability and filtering

The posterior distribution, i.e., the probability of each gene *j* being ‘truly expressed’ in cell *i* is given by 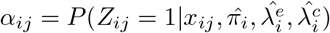, where:

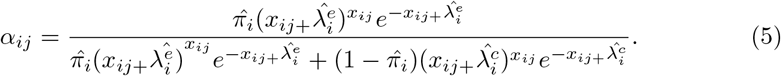

Here 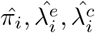 denote their respective MLE estimates derived from the above EM algorithm. A gene is filtered (i.e., count set to 0) if *α*_*j*_ ≤ 0.75. Although 0.75 is the default cutoff value, it is a tunable parameter by the user. To determine this default value, empirical analysis was conducted on benchmark datasets (Supplementary Note 4).

It should be noted that the DenoIST model assumes that gene filtering is all-or-nothing, i.e., a positive gene count is either contamination or endogenous expression, with no in-between. This assumption likely only works for small targeted panels like current IST data and should be reconsidered before applying to whole transcriptome measurements. For more discussion on this assumption please refer to Section 3.

### 5.5 Alternative segmentation and count correction methods

#### 5.5.1. Default

For Xenium data, Xenium Ranger (v3.1.1) was run with default parameters. The default segmentation refers to the boundary expansion approach implemented by Xenium Ranger, which is obtained by running “import-segmentation” with the option “–expansion-distance=15”. For MERSCOPE and CosMx data, the downloaded count matrices were used directly.

### 5.6 Nuclei transcripts only

To restrict the transcripts to nuclei transcripts only, we filtered the transcript files to keep only transcripts overlapping the nucleus. For Xenium and CosMx data, we kept transcripts being originally labelled as ‘overlapping nucleus’ for Xenium or being in the ‘Nuclear’ compartment in CosMx. For MERSCOPE data, as this information was not readily available, we kept transcripts that are less than 5 µm away from the cell centroid. Then, ‘tx2spe()’ function from SubcellularSpatialData (v1.6.0) was used to convert the transcript files to count matrices.

#### 5.6.1 Proseg

Proseg (v3.0.10) was run with default parameters. Proseg takes the transcript file as input, and the “–xenium” preset was used for all Xenium datasets. The “–merscope” and “–cosmx” presets were used for MERSCOPE and CosMx data respectively. The output count matrices were directly used.

#### 5.6.2 ResolVI

ResolVI (scvi-tools v1.4.0.post1) was run with default parameters. The model was trained in an unsupervised manner for 100 epochs as recommended by the tutorial. Counts are generated according to the tutorial at https://docs.scvi-tools.org/en/stable/tutorials/notebooks/spatial/resolVI_tutorial.html (Date accessed: 5/6/2025). Briefly, the median of posterior was used with options “num samples=3” and “summary frequency=30”.

#### 5.6.3 SPLIT

SPLIT (v0.1.2.1) was run following the tutorial at https://github.com/bdsc-tds/SPLIT/blob/main/vignettes/Run RCTD and SPLIT on Xenium.Rmd (Date accessed: 23/6/2025). RCTD (spacexr v2.2.1) was first run followed by SPLIT with UMI min = 1 to avoid over filtering cells.

### 5.7 Evaluation of noise removal

To evaluate the performance of tools, we used the following metrics.

#### 5.7.1 Mutually Exclusive Co-Expression Rate

The Mutually Exclusive Co-Expression Rate (MECR) metric was originally proposed by Hartman and Satija [19], and has been adopted in a few recent pre-prints as well [6, 13]. Briefly, for a given pair of genes *g*_1_ and *g*_2_, the MECR can be calculated by:

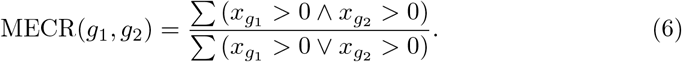

The pairs of genes that are assumed to be mutually exclusive are selected from reference scRNA-seq datasets for each corresponding tissue or cell type. Genes with high specificity to a cell type are selected by first performing differential expression (DE) analysis on the reference dataset with ‘scran’ (v1.36.0) Lun et al. [24], then filtering for DE genes that are: 1) top 30% by mean AUC, 2) detected in ≥ 50% of a cell type, and 3) detected in ≤ 5% of the rest of the cells. The resulting genes are then paired up to form mutually exclusive pairs. This yielded 337, 45, 44, 266, and 110 mutually exclusive gene pairs for the Xenium breast cancer, Xenium healthy lung, Xenium mouse brain, CosMx mouse brain, and MERSCOPE mouse brain datasets respectively.

For resolVI and SPLIT, as they do not return integer counts, counts are rounded to the nearest integer before calculating MECR, i.e. *x* ≥ 0.5 is considered to be ‘positive’. Note that this does not affect values larger than 0.5 as MECR only considers non-zeroes vs zeroes.

#### 5.7.2 Positive Marker Purity

The Positive Marker Purity (PMP) metric is proposed by Heidari et al. [6] as a true positive metric. It measures the proportion of cell-type marker transcripts in each cell. Formally, for each cell *i* and a set of pre-defined marker genes *M*, the PMP is defined as:

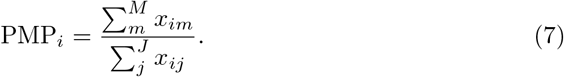

Cell-type markers are defined by performing differential expression (DE) testing on the corresponding scRNA-seq reference data. DE testing is done with the scoreMarkers function from the R package ‘scran’ (v1.36.0) [24], using rank.logFC.cohen ≤ 5 as the cutoff. The cell type of each cell is determined with label transfer from the reference scRNA-seq data using SingleR (v2.10.0) [25]. The mean PMP is calculated by taking the mean within each cell type first, followed by taking the mean of all cell types.

#### 5.7.3 RCTD analysis

RCTD (spacexr v2.2.1) was run on the healthy lung sample VUHD116A before and after denoising. The default settings were used with UMI min = 1 to avoid over filtering of cells, and ‘run.RCTD’ was run with ‘doublet’ mode.

## Supplementary information

Supplementary Figures S1–S3, Supplementary Notes 1–4.

## Acknowledgements

The authors acknowledge Ruqian Lyu, Saahithi Mallapragada and Angela Oill for helpful discussions about this manuscript and testing of initial versions of the software.

## Declarations

### Funding

This work was supported by US National Institutes of Health grant R01HG011886 awarded to N.E.B., J.A.K., and D.J.M., R01HL145372 awarded to N.E.B., J.A.K., and Australian National Health and Medical Research Council grant GNT1195595 awarded to D.J.M.

### Competing interests

The authors declare no competing interests.

### Ethics approval and consent to participate

Not applicable.

### Consent for publication

Not applicable.

### Data availability

Data and code to reproduce the figures in this manuscript are available (under CC BY 4.0 license) at the following Gitlab repository: https://gitlab.svi.edu.au/ biocellgen-public/deno 2025 denoist paper reproducibility and Zenodo (https://doi.org/10.5281/zenodo.17595028). Accessions for third-party and publicly available datasets used are listed in Table 1.

### Code availability

DenoIST is available on Github (https://github.com/aaronkwc/DenoIST) as an R package. DenoIST takes an IST count matrix as input and returns an adjusted count matrix with contamination removed. It is designed to work with the SpatialExperiment class from Bioconductor, but is also compatible with normal matrix and dataframe input types. Parallelisation over CPUs is also available to speed up computation. A vignette demonstrating its usage on Xenium data is accessible from the Github repository.

### Author contribution

D.J.M., N.E.B., J.A.K., and A.V. identified the issue and conceptualised the study. D.J.M and H.S. supervised the study. A.K. developed the methodology, implemented the package, performed all data analysis and prepared all figures. D.J.M, H.S., A.K. wrote the manuscript. A.V., N.E.B, J.A.K, reviewed and provided feedback on the package and manuscript. All authors reviewed the manuscript. D.J.M., N.E.B., J.A.K. secured funding.

## Notes

https://gitlab.svi.edu.au/biocellgen-public/deno_2025_denoist_paper_reproducibility

### Competing Interest Statement

The authors have declared no competing interest.

