## Supplementary Information for "Denoising image-based spatial transcriptomics data with DenoIST"

### Supplementary note

November 2025

4 Empirical analysis on posterior threshold value 7

#### List of Figures

S1 Density plot of non-zero gene counts . . . . . 4
S2 Empirical analysis on posterior thresholds . . . . . 7
S3 Relationship between mean MECR and mean PMP with changing posterior threshold . . 8

#### 1 Data generative proccess for IST data

We work with an $N \times G$ 'count matrix' X which holds information about the number of observed counts in N cells and G genes. We let $i \in \{1, \ldots, N\}$ index cells and $j \in \{1, \ldots J\}$ index genes, so that $x_{ij}$ is the observed count of the jth gene in the ith cell. Let the true number of molecules (unobserved) of gene j in cell i be $m_{ij}$. We follow the nomenclature defined by Sarkar and Stephens [1] and define our model with a measurement and expression component.

##### 1.1 Spatial measurement model

The measurement model of scRNA-seq comes from the assumption of having a finite number of reads in each cell distributed among G genes. This results in the multinomial model for each cell:

$$x_{i1} \ldots x_{iJ} \mid x_{i+}, \lambda_{i1} \ldots \lambda_{iJ} \sim \text{Multinomial}(x_{i+}; \lambda_{i1} \ldots \lambda_{iJ}), \tag{1}$$

where $\lambda_{ij} = m_{ij}/m_{i+}$ is the relative expression of gene j in cell i with respect to the cell itself, and $x_{i+} = \sum_j x_{ij}$ is the total detected transcript count of cell i.

However, during the measurement process in image-based spatial transcriptomics data, the repeated unit of measurement is not a cell, as they are artificial boundaries added post-hoc, via cell segmentation. Instead, the repeated unit measurement is each gene or each gene probe. For each gene, a finite number of detectable probes is distributed among N cells or any cell-like objects with arbitrary boundaries. This conception of the process results in a transposed version of Eq. 1:

$$x_{1j} \dots x_{Nj} \mid x_{+j}, \lambda_{1j} \dots \lambda_{Nj} \sim \text{Multinomial}(x_{+j}; \lambda_{1j} \dots \lambda_{Nj}), \tag{2}$$

where $\lambda_{ij} = m_{ij}/m_{+j}$ is that the relative expression of gene j in cell i with respect to the whole population of cells, and $x_{+j} = \sum_i x_{ij}$ is the total detected transcript count of gene j. A key assumption here is the binding and detection of probes is independent between genes.

A well-known result is that the multinomial likelihood can be simplified to a Poisson likelihood, under the assumption that the true total number of transcripts is much greater than the total number of molecules detected, i.e., $m_{ij} << m_{+j}$:

$$x_{ij} \mid x_{+j}, \lambda_{ij} \sim \text{Poisson}(x_{+j}\lambda_{ij}). \tag{3}$$

Essentially, this is a measurement model per cell with gene total count as the offset, which is a transposed version of conventional scRNA-seq methods.

To incorporate spatial information into the measurement model, we encoded it into the offset $x_{+j}$ by only summing counts in neighbouring cells. This is based on the intuition that contamination is likely a more local effect that involves cells in close proximity. Hence, each cell has its specific offset based on its neighbours.

For each cell i, the offset $x_{ij+}$ is calculated by:

$$x_{ij+} = x_j^b + \sum_{t' \in T} x_{t'j}, \tag{4}$$

where T denotes the set of neighbouring cells to cell i. The default option to define T is to include cells with centroids within 50 µm from the centroid of cell i. A distance based inclusion is to account for neighbourhood densities, as denser tissues tend to exhibit more contamination due to cells being more closely packed together and segmentation difficulty is higher. However this distance d is tunable by the user.

To account for global ambient contamination on top of local neighbourhood effects, a background offset $x_j^b$ that is shared globally among all cells can be added to $x_{ij+}$. To calculate $x_j^b$, transcripts are binned into 200 hexbins by default. To determine which spatial bins represent background (i.e., areas not primarily occupied by cells), a Gaussian Mixture Model (GMM) is used. The total transcript counts across all spatial bins are assumed to follow a 2-component mixture of Gaussian distributions: one representing background with low transcript counts and the other corresponding to bins with biologically meaningful signal, i.e., higher transcript counts.

Formally, let $C_k$ denote the total transcript count in hexbin k. The GMM models the density of $C_k$ as:

$$P(C_k) = w \cdot \mathcal{N}(C_k \mid \mu_1, \sigma_1^2) + (1 - w) \cdot \mathcal{N}(C_k \mid \mu_2, \sigma_2^2), \tag{5}$$

where w denotes the proportion of background bins, $\mu$ and $\sigma^2$ are the mean and variance of their respective components. The GMM is fit using the Expectation Maximisation (EM) algorithm, as implemented in the 'flexmix' R package. After model fitting, each hexbin is assigned to the most likely component based on its posterior probability. The component with the lower $\mu$ is designated as the background component. The global background expression level for gene j, denoted $x_j^b$, is then computed by averaging the gene's expression values across all bins classified as background, and then scaled to the local neighbourhood size defined by the distance d:

$$x_j^b = \frac{\pi d^2}{A} \frac{1}{|\mathcal{B}|} \sum_{i \in \mathcal{B}} x_{ij}, \tag{6}$$

where $\mathcal{B}$ denotes the set of bins assigned to the background component, and A denotes the area of one hexbin which can be calculated with basic geometry, based on the size of the input sample and number of bins used.

This global background profile is subsequently incorporated into the DenoIST model as an additive offset to account for both local contamination and global ambient signal.

##### 1.2 Expression-Contamination model

To incorporate the contamination effect into the measurement model, we further specify the mean parameter $\lambda_{ij}$ to have two distinct values for each cell, such that the counts $x_{ij}$ can either come from true expression from cell i or contamination from neighbouring cells. Combining with Eq. 3 results in a Poisson mixture model:

$$x_{ij} \sim \pi_i \text{Poisson}(x_{ij+} \lambda_i^e) + (1 - \pi_i) \text{Poisson}(x_{ij+} \lambda_i^c), \tag{7}$$

where $\lambda_i^e$ denotes the mean parameter of genes that are truly expressed in cell i and $\lambda_i^c$ denotes the mean parameter of genes that are contaminated in cell i.

Note that it is a transposed version of conventional gene-wise models used in scRNA-seq, with spatial information encoded within the offset term.

#### 2 Motivations for the DenoIST mixture model

The motivation for the per-cell mixture model came from an observation from the Xenium healthy lung dataset. In this dataset, even though only nuclei transcripts were kept, the annotated immune cells often present higher transcript counts for epithelial and endothelial marker genes than for immune markers. However, if the local neighbourhood is factored in, the proportion of immune marker gene counts relative to the neighbourhood is actually high, i.e., compared to cells in close proximity, the annotated immune cells express relatively high counts for immune gene markers, while the non-immune markers are relatively lowly expressed.

To better illustrate the above motivation, here I show an example of a single annotated B cell in the Xenium healthy lung dataset (Fig. S1). The raw gene counts were divided by the total count of the

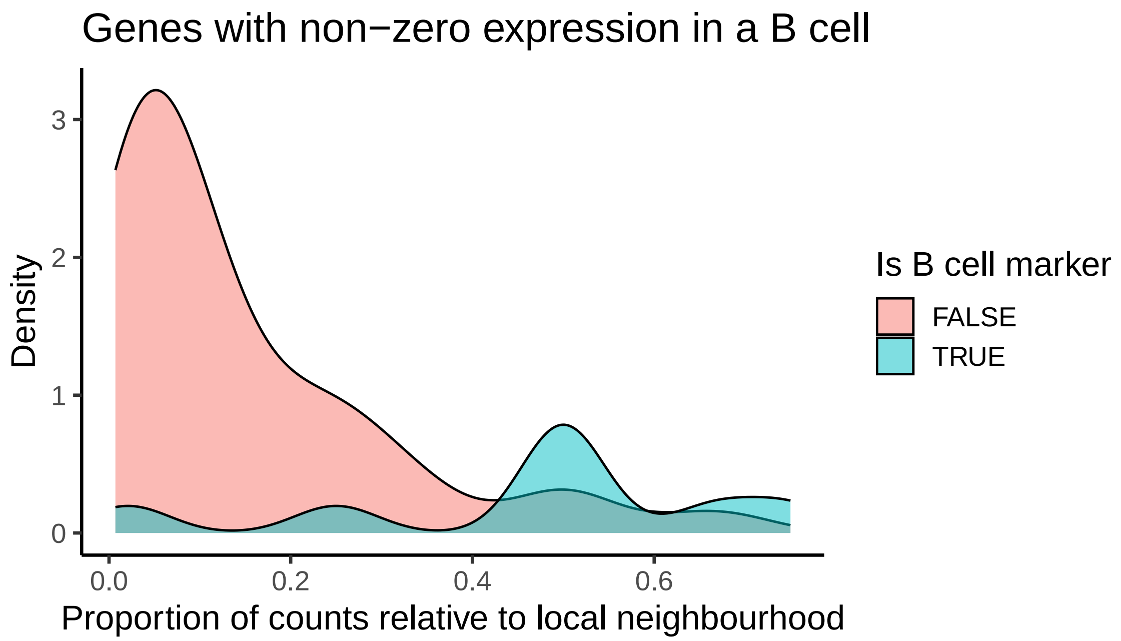

Figure S1: Density plot of non-zero gene counts in an annotated B cell from the Xenium healthy lung dataset, proportionate to the cell's local neighbourhood counts. B cell marker genes used here are *CD19*, *CD79A*, *CD79B*, *MS4A1*, *BANK1*, *TCL1A*, *TNFRSF13C*, and *PTPRC*.

cell's neighbourhood of radius of 50 µm. Even though the raw count of B cell marker *MS4A1* is less than that of endothelial marker *EPAS1* ($x_{MS4A1} = 3, x_{EPAS1} = 6$), the total count in the neighbourhood shows a large difference ($x_{MS4A1+} = 4, x_{EPAS1+} = 122$). Thus, the proportion of counts relative to local neighbourhood for *EPAS1* is substantially lower than that of *MS4A1*, which is consistent with the clustering and annotation. Considering all the genes with non-zero expression, the count proportions manifest a mixture distribution (Fig. S1), further motivating the use of a per-cell mixture model.

#### 3 Derivation for expectation maximisation (EM) algorithm

For brevity, we will drop the i index in the following discussion as parameters are estimated for each cell separately. Let $\theta = \{\pi, \lambda^e, \lambda^c\}$ be the set of parameters needing to be estimated from data. Let $Z_j$ be the latent identity of whether gene j is truly expressed, such that:

$$Z_j \sim \text{Bernoulli}(\pi). \tag{8}$$

Then the complete data log likelihood is:

$$\begin{aligned} \log P(Z, X|\theta) = \sum_j [&Z_j \log p(x_j|\lambda^e) + (1 - Z_j) \log p(x_j|\lambda^c) \\ &+ Z_j \log \pi + (1 - Z_j) \log(1 - \pi)]. \end{aligned} \tag{9}$$

The Q function is the expectation of the complete log likelihood w.r.t. $Z_j|X, \hat{\theta}$, which is:

$$Q(\theta, \hat{\theta}) = \sum_j [\gamma_j \log p(x_j|x_{j+}, \lambda^e) + (1-\gamma_j) \log p(x_j|x_{j+}, \lambda^c) \\ + \gamma_j \log \pi + (1-\gamma_j) \log(1-\pi)]. \tag{10}$$

#### 3.1 E-step

For the E-step, I initialise some values to find the expected complete data log likelihood.

$$\begin{aligned}
\gamma_j &= \mathbb{E}[Z_j|x_j, \hat{\theta}] \\
&= P(Z_j = 1|x_j, \hat{\theta}) \\
&= \frac{\pi \frac{(x_{j+}\lambda^e)^{x_j} e^{-\lambda^e}}{x_j!}}{\pi \frac{(x_{j+}\lambda^e)^{x_j} e^{-x_{j+}\lambda^e}}{x_j!} + (1-\pi) \frac{(x_{j+}\lambda^c)^{x_j} e^{-x_{j+}\lambda^c}}{x_j!}} \\
&= \frac{\pi (x_{j+}\lambda^e)^{x_j} e^{-x_{j+}\lambda^e}}{\pi (x_{j+}\lambda^e)^{x_j} e^{-x_{j+}\lambda^e} + (1-\pi)(x_{j+}\lambda^c)^{x_j} e^{-x_{j+}\lambda^c}}.
\end{aligned}$$

#### 3.2 M-step

For the M-step, closed form solutions can be derived with partial differentiation.

For $\pi$, partially differentiate Eq. 10 w.r.t $\pi$ and equate to 0:

$$\frac{\partial}{\partial \pi} Q(\theta, \hat{\theta}) = 0$$

$$\sum_{j}^{G} [\gamma_j (\frac{1}{\pi}) + (1 - \gamma_j) \frac{-1}{1 - \pi}] = 0$$

$$(\frac{1}{\pi}) \sum_{j}^{G} \gamma_j + \frac{-1}{1 - \pi} \sum_{j}^{G} (1 - \gamma_j) = 0$$

$$(\frac{1}{\pi}) \sum_{j}^{G} \gamma_j + \frac{\sum_j^G \gamma_j - G}{1 - \pi} = 0$$

$$\sum_{j}^{G} \gamma_j (\frac{1}{1 - \pi} + \frac{1}{\pi}) - \frac{G}{1 - \pi} = 0$$

$$\frac{G}{1 - \pi} = \sum_{j}^{G} \gamma_j (\frac{1}{1 - \pi} + \frac{1}{\pi})$$

$$\frac{G}{1 - \pi} = \sum_{j}^{G} \gamma_j \frac{(\pi + (1 - \pi))}{(1 - \pi)\pi}$$

$$\pi = \frac{\sum_j^G \gamma_j}{G} = \bar{\gamma}_j.$$

For $\lambda^e$, partially differentiate Eq. 10 w.r.t $\lambda^e$ and equate to 0:

$$\frac{\partial}{\partial \lambda^e} Q(\theta, \hat{\theta}) = 0$$

$$\frac{\partial}{\partial \lambda^e} \sum_{j} \gamma_j \log( \frac{(x_{j+} \lambda^e)^{x_j} e^{-x_{j+} \lambda^e}}{x_j!} ) = 0$$

$$\frac{\partial}{\partial \lambda^e} \sum_{j} \gamma_j [x_j \log(x_{j+}) + x_j \log(\lambda^e) - x_{j+} \lambda^e - \log(x_j!)] = 0$$

$$\frac{\partial}{\partial \lambda^e} \sum_{j} \gamma_j [x_j \log(\lambda^e) - x_{j+} \lambda^e] = 0$$

$$\frac{\sum \gamma_j x_j}{\lambda^e} - \sum_{j} x_{j+} \gamma_j = 0$$

$$\lambda^e = \frac{\sum_j \gamma_j x_j}{\sum_j \gamma_j x_{j+}}.$$

For $\lambda^c$, the same derivation yields the closed form:

$$\lambda^c = \frac{\sum_j (1 - \gamma_j) x_j}{\sum_j (1 - \gamma_j) x_{j+}}.$$

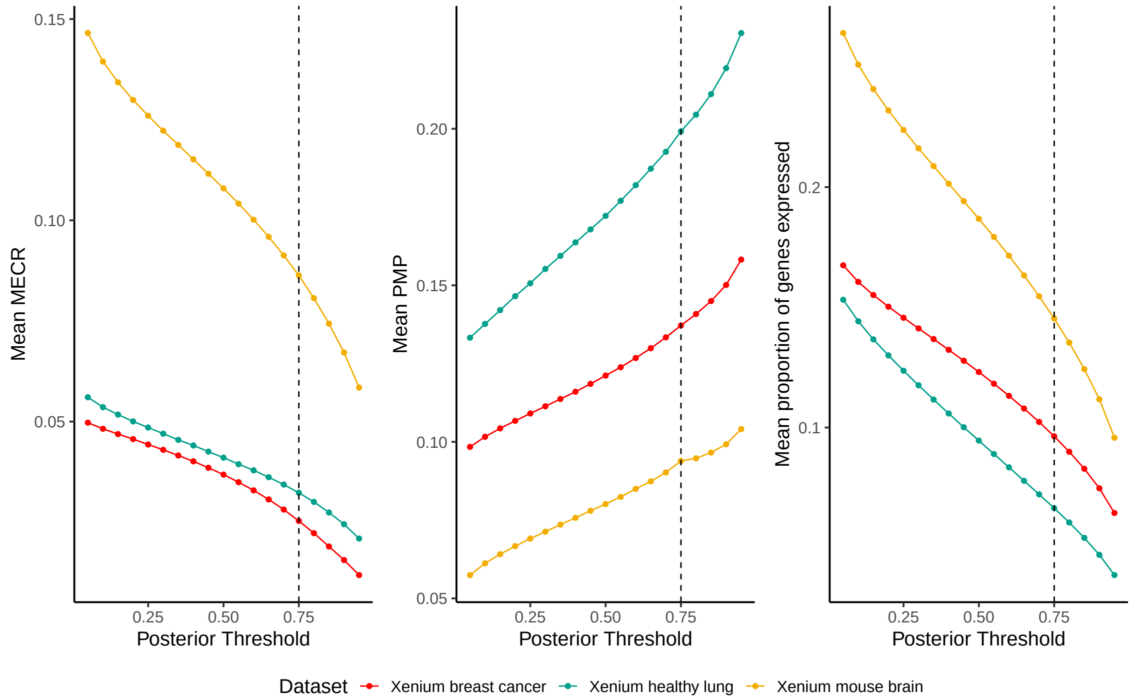

Figure S2: Empirical analysis on how benchmarking metrics vary with posterior threshold. Metrics shown here are (Left) mean MECR, (Middle) mean PMP, and (Right) mean proportion of genes expressed. Dotted line indicates default value provided in DenoIST.

### 4 Empirical analysis on posterior threshold value

DenoIST provides a default posterior cutoff value of 0.75, i.e., genes with $\alpha \leq 0.75$ are filtered. This threshold value is empirically determined using the benchmarking datasets with boundary expansion segmentation, by comparing the mean MECR, mean PMP, and mean proportion of genes expressed over a range of thresholds (Fig. S2). Note that thresholds at 0 and 1 correspond to no filtering and all counts filtered respectively.

In general, all metrics monotonically increase or decrease with the threshold, e.g., the mean MECR (false positive rate) decreases as the filtering becomes more stringent, while the mean PMP (true positive rate) increases (Fig. S3). That behaviour means DenoIST would benefit in benchmark performances if the threshold is as stringent as possible as the metrics would be the most optimal. However, this performance with respect to the metrics comes at the cost of reducing the number of genes detected. Limiting detection rate naturally decreases the likelihood of erroneous assignments, but it also implies that true biology is more likely to be missed. To err on the side of caution, a threshold of 0.75 was chosen to maintain a 10-15% detection rate on average, but it is a tunable parameter by users depending on the desired trade-off. The results observed from varying the posterior threshold used for filtering indicate the theoretical cap in benchmark performance of DenoIST (Fig. S3).

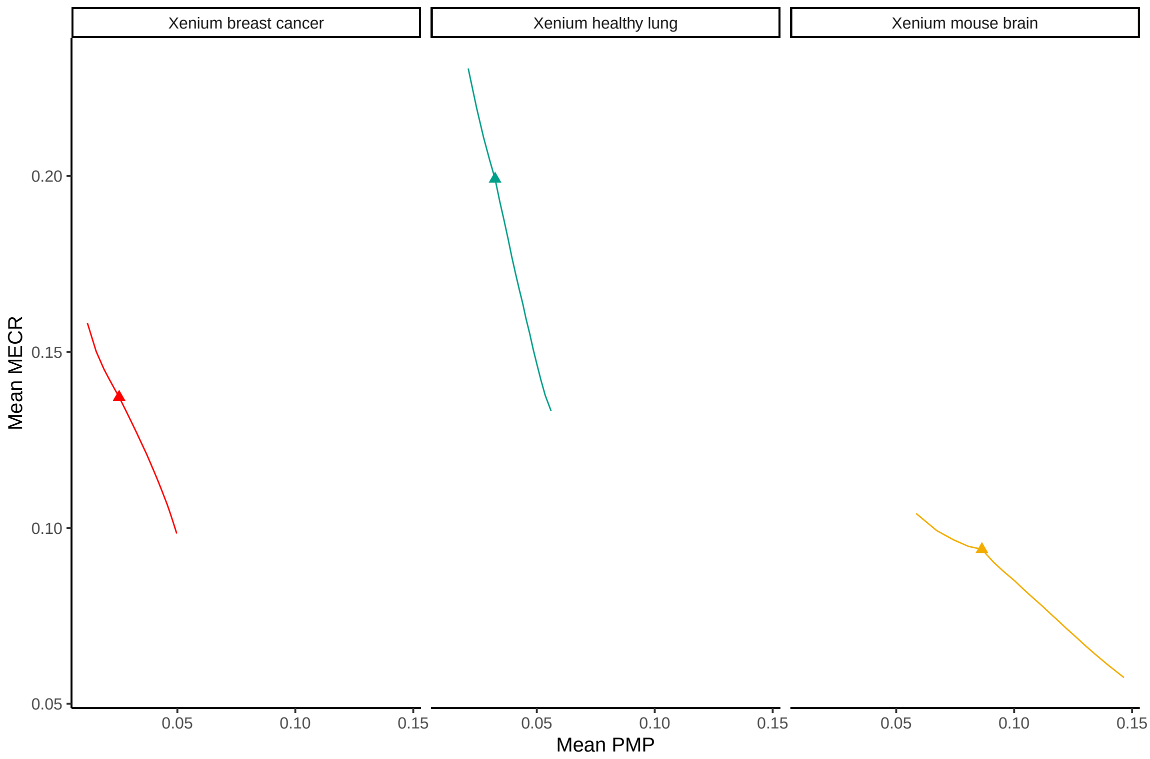

Figure S3: The relationship between mean MECR and mean PMP when the posterior threshold changes. Highlighted point indicates the default threshold value (0.75).
